# Beta-tACS does not impact the dynamics of motor memory consolidation

**DOI:** 10.1101/2020.05.25.114850

**Authors:** Liliia Roshchupkina, Whitney Stee, Philippe Peigneux

## Abstract

The consolidation of motor memory is a non-linear temporal dynamic. There are critical time points at which post-training performance can improve (e.g., 30 min and 24 h) or merely stabilize (e.g., 4 h). Besides, neuronal plasticity is supported by synchronized oscillatory activity in and between brain areas at play during the acquisition and consolidation of motor skills. Transcranial alternating current stimulation (tACS) can entrain cortical oscillatory activity, which may eventually modulate brain plasticity-related processes. Previous reports suggest that 20 Hz electrical stimulation over the primary motor cortex (M1) following training facilitates the consolidation of motor memories. To the best of our knowledge, the effect of tACS was not investigated when applied at critical post-training time points, nor its impact at longer time scales. In the present study, we investigated the effect of 20 Hz tACS applied over M1 within critical time periods (25min vs. 4h) on motor memory consolidation at immediate and delayed (24h) retrieval. Performance similarly evolved over time in all conditions, independently from the stimulation type (20 Hz tACS vs. Sham tACS) or the stimulation time point. As it stands, our results do not support the proposal that 20 Hz tACS exerts a positive, specific effect on the consolidation of motor memories.

## Introduction

The fine-tuned motor activities that are at the core of our everyday life require skilled coordination. Even a simple motor task such as preparing one’s morning coffee requires learning to integrate basic motor actions into a coherent sequence, until it becomes a routine. Indeed, a motor skill is not instantaneously created. Rather, it develops over successive steps both during actual practice and post-training time. In a fast phase, performance rapidly increases online with motor practice, eventually followed by slower gains developing offline (i.e., without actual rehearsal) during post-practice time [1]. Fast and slow processes are thought to correspond to synaptic and system consolidation, respectively [2]. Furthermore, the offline consolidation of recently learned motor skills is a multi-step, dynamic process. Post-training motor performance is usually markedly increased in a “boost” phase taking place within the first 5 to 30 minutes after the end of practice [2-6] as well as 24h to 48h later [2-4]. However, performance remains at the level reached at the end of the learning session when tested in a “silent” period 4 to 5 hours after [2-4]. Additionally, the early boost in performance was found predictive of performance levels eventually achieved 48h later [2], suggesting the functional relevance of immediate post-training periods in the development of longer-term memory consolidation processes.

Motor acquisition and early consolidation processes are associated with task-induced oscillatory changes over the primary motor cortex (M1) [7, 8], specifically in the beta (β) band (13–30 Hz) [9]. Consequently, researchers aimed at manipulating motor learning and memory using transcranial alternating current stimulation (tACS) that allows delivering rhythmic electrical stimulations over the scalp. Indeed, the effects of tACS are primarily attributed to the entrainment of the endogenous oscillatory brain activity to the stimulation frequency [10, 11]. As well, it was proposed that tACS-related entrainment of echoes and spike-timing-dependent plasticity [12] triggers local excitability and connectivity changes during the hour post-stimulation [13], [14]. Accordingly, tACS was shown to modulate local motor cortical excitability and long-range functional connectivity in a frequency-dependent manner [12, 13]. At the behavioural level, 20 Hz tACS over M1 was found to facilitate the acquisition [17] and retrieval [18] of a motor sequence. In the latter study, reaction times markedly improved at retrieval immediately after 20 Hz tACS, as compared to other conditions (i.e., 10-Hz tACS, sham stimulation or effective transcranial direct current stimulation [tDCS]). This led to the hypothesis that 20 Hz tACS increases motor-cortical excitability and neuroplasticity, which in turn promotes early motor memory consolidation. On the other hand, no behavioural effects were found when retesting the motor sequence 6h after β-tACS over M1 [19]. This lack of behavioural effect might be due to the fact that delayed retrieval performance was assessed during the silent period mentioned above, during which spontaneous performance gains are usually not observed [2]. Accordingly, repetitive low frequency (1Hz) transcranial magnetic stimulation (rTMS) M1 immediately after learning was found to abolish the 30-minutes performance boost while leaving unimpaired delayed behavioural improvement at 24 or 48h [3]. It indicates that, on the one hand, the short-term performance boost reflects to some extent the motor system capabilities for further offline improvements, and, on the other hand, that long-term consolidation processes do not solely depend on neural optimization in M1 and engage additional cortico-subcortical circuits.

To the best of our knowledge, the effect of electrical cortical stimulations during critical post-training time windows on the evolution of motor memory consolidation has not been systemically tested. Here, we aimed at probing the effect of 20 Hz tACS on M1 on the evolution of motor performance at 25 min (boost), 4h (silent) and 24h post-training windows. We hypothesized (1) that 20 Hz tACS delivered immediately after the end of learning (early boost) would modulate motor learning-related neuronal networks activity and promote higher performance gains; (2) that a repeated 20 Hz tACS at the end of the boost period would further increase these effects; and (3) that 20 Hz tACS administered within the silent period immediately before retesting could optimize the motor system for task execution and enhance performance as well as promote further long-lasting changes.

## Material and methods

All volunteers gave written informed consent to participate in this study approved by the ULB-Erasme Ethics Committee (approval P2018/480). In total, 73 healthy right-handed participants aged 18-30 years were recruited. Only subjects with a moderate to neutral chronotype score (between 30 and 70 at the Morningness-Eveningness Questionnaire [20]) and good sleep quality (score ≤ 7 at the Pittsburgh Sleep Quality Index [21]) were included. Computer scientists and musicians who already exhibited high-level hand dexterity, smokers, and individuals with neurological/psychiatric medical records or exposed to jetlag within the past 3 months were excluded.

### Experimental procedure

All participants followed a similar behavioural procedure with one motor learning session and 3 retest sessions at three specific time points post-learning: 25min, 4h, and 24h. They were randomly assigned to one out of the 5 possible Group conditions differing by the timing of tACS administration (immediate post-learning, 25min or 4h) and type (Sham vs. Actual tACS; see Fig. 1). Female participants were tested within the second week of their menstrual cycle to avoid a potential memory consolidation bias induced by changes in hormonal levels [19, 20].

**Fig 1.**
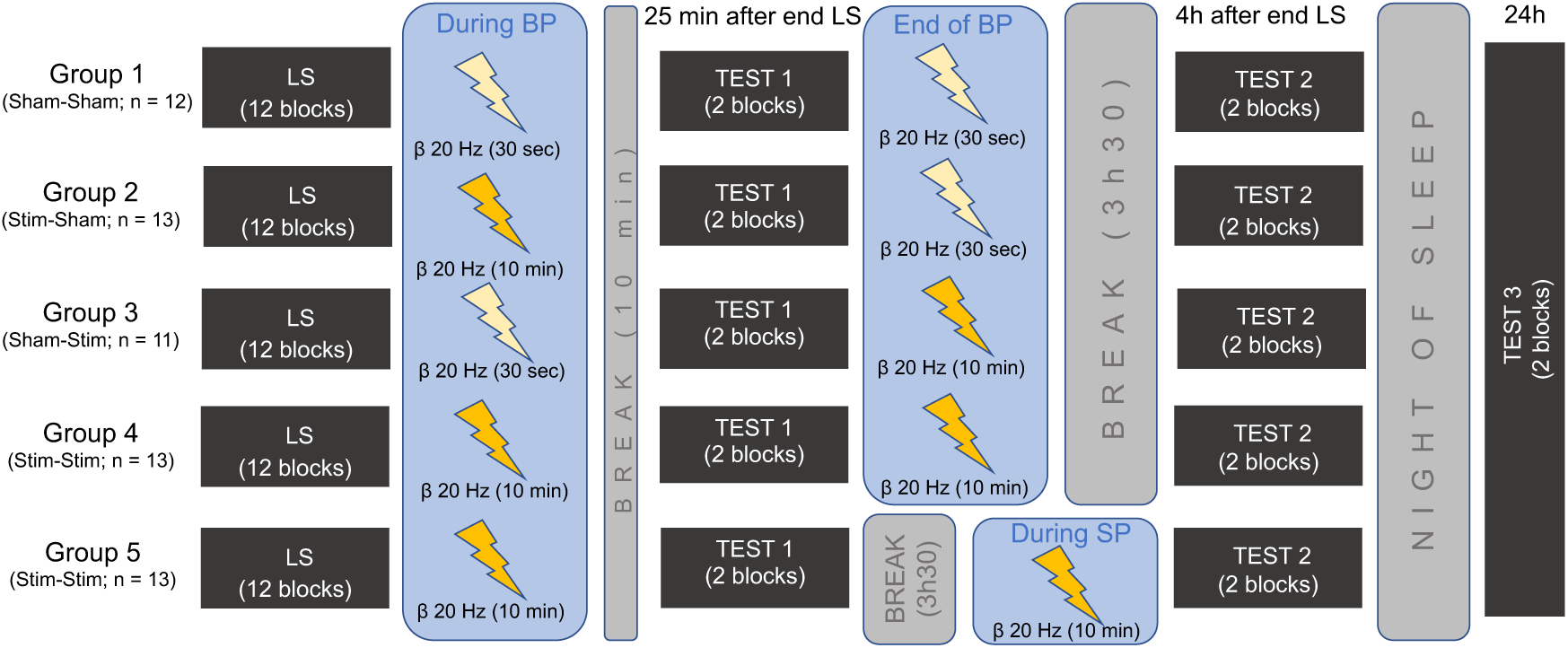
Experimental design: All participants were trained on 12 FTT blocks during the learning session (LS). Learning was immediately followed by a 10-minutes stimulation period during which participants were administered either a Stimulation (10 minutes 20 Hz tACS) or a Sham (30-seconds 20 Hz tACS only at the onset of the 10-minutes period) condition. After a ±10 minutes break (i.e., 25 minutes after LS), motor performance was assessed over 2 FTT blocks (Test 1), immediately followed again by a Sham or Stimulation (Group 1-4) condition at the end of the boost period (BP), or no stimulation (Group 5). All participants came back to the lab 4 hours after the end of LS to be tested again over 2 FTT blocks (Test 2) during the silent period (SP). Participants from Group 5 were administered a Stimulation condition immediately before testing. Approximately 24 hours after the end of LS, volunteers came back to perform the last testing session over 2 FTT blocks (Test 3).

### Motor learning task (FTT)

The experiment was conducted using Cogent 2000 (http://www.vislab.ucl.ac.uk/cogent.php) running on Matlab 6.1. (Mathworks, USA). Participants were trained on a 5-elements Finger Tapping Task (FTT) adapted from [1]. They were explicitly asked to reproduce as fast and accurately as possible a 5-elements sequence of finger movements for 30 seconds (i.e., one block) using their non-dominant hand. Each finger corresponded to one digit (from 1 = little finger to 4 = index), and the sequence to reproduce (4-1-3-2-4) was on permanent display on the computer screen during execution. The learning session (LS) consisted of 12 blocks separated by 20-second rest periods. Immediately after the end of the learning phase, the first stimulation (tACS) session was administered for 10 minutes, either using an actual Stimulation (20 Hz tACS for 10 minutes; Groups 2, 4, 5) or a Sham (20 Hz tACS only for the first 30 seconds of the 10-min period; Groups 1, 3) condition, pseudo-randomly counterbalanced (see Fig.1). Twenty-five minutes after the end of the learning session (i.e., +/-10 minutes after the end of the first stimulation session), all participants were administered 2 FTT blocks (Test 1) to assess motor performance within the boost period. Immediately after, they received a second 10-minutes tACS under a Stimulation (Groups 3, 4) or Sham (Groups 1, 2) condition, or could leave the laboratory without stimulation (Group 5). Participants came back to the laboratory 4 hours after the end of the learning session to be tested again on 2 blocks (Test 2) during the so-called silent period. In Group 5, the Stimulation condition (10-minutes 20 Hz tACS) was administered immediately before behavioural testing. Finally, participants came back the next day at the same time as the learning session took place (i.e., after 24h) to be tested for the third time on 2 FTT blocks (Test 3).

### TACS parameters

Transcranial alternating current stimulation (tACS) was delivered using a DC-Stimulator Plus (NeuroConn GmbH, Ilmenau, Germany) device at an intensity of 1 mA peak to peak, which corresponds to a current density of 0.04 mA/cm^2^ under the pads. The sinusoidal current was applied at a frequency of 20 Hz over two conductive rubber pads (5cmx5cm) pasted using Ten20^TM^ conductive paste. The active pad was centred over the right M1, and the reference pad was located on the contralateral deltoid. We located the right M1 area using the International 10-20 System [24] where it matches the C4 location [25], 20% to the right of the Cz location. The stimulator was programmed to ramp up/down within 10 seconds at the beginning/end of the stimulation period. After each period of stimulation, participants were asked to report the strength of the potential sensory sensations (auditory, visual, cutaneous and gustatory) they perceived during stimulation on a scale ranging from 0 to 10.

### Additional measures

To ensure that all groups benefitted from the same amount of sleep before and during the procedure, sleep duration and sleep-wake cycle regularity were assessed for the 3 preceding nights as well as for the night following the learning session, using a self-reported sleep questionnaire [26] and actimetry (wGT3X-BT; Actigraph, USA). Also, before the learning session, Test 2 and Test 3, they were administered a 5-minutes version of the Psychomotor Vigilance Test (PVT5 [27]). PVT performance was evaluated using the reciprocal reaction time (RRT = 1/RT) shown to be most sensitive to detect changes in vigilance [28]. They also completed the Visual Analogue Scale for Sleepiness and Fatigue (adapted from [29]) 8 times, before and after the learning session and before and after each test session.

### Data analysis

FTT performance was analysed computing the General Performance Index (GPI) that takes into account both speed and accuracy [30]. For the learning session, the GPI was considered for each block separately. For testing sessions and comparison with performance achieved at the end of the learning session, a mean GPI was computed over the two blocks of each test (Tests 1-3) as well as the last two blocks of the learning session. For the purpose of comparisons with prior reports, similar analyses were computed using the speed parameter only (i.e., mean RT per block for correct responses; see Supplementary Information [SI] results).

Mixed-design analyses of variances (ANOVAs) were computed on SPSS (IBM SPSS Statistics version 24) with between subjects’ factor Group (1-5) and within-subject factor Session (end of LS, Test 1-3) or Block (within the learning session). Sphericity assumption was tested using Mauchly’s test. When needed, degrees of freedom were corrected using Greenhouse-Geisser sphericity estimates. Post-hoc pairwise comparisons in the presence of significant interaction effects were conducted with Bonferroni’s adjustment for multiple comparisons.

For demographical and tACS impedance data, Welch’s ANOVAs were run with between subjects’ factor *Group* (1 to 5). Welch’s ANOVAs were preferred to Classical Oneway ANOVAs considering their increased power in case of heterogeneity of variances, that Levene’s Test for equality of variances often fails to detect [31]. For stimulation data, Mann-Whitney’s tests were run with between-subject factor *Condition* (Sham vs. Actual stimulation).

## Results

Out of the 73 participants, 10 were excluded due to excessive tACS impedance (> 10 kOhm), 1 due to poor accuracy at different sessions (< 80%). Eventually, 62 subjects (43 females; mean age = 21.47 ± 2.30 yrs, min = 18, max = 29) were included. Age, laterality, circadian rhythm, and sleep quality were not significantly different between the 5 groups (Welch’s ANOVAs, *p*s > 0.285; see SI Table 1). Likewise, alertness, sleepiness, fatigue, sleep, stimulation parameters and reported stimulation sensations were not different between conditions or modulated by the Group condition (all *p*s > 0.117; see SI analyses).

Sleep duration was similar between the 5 Group conditions for the 3 nights prior to the experiment (average 8.23 ± 1.31 hours; min = 4h, max = 12h; Mixed ANOVA *F*_(2, 8)_ = 11.859, *p* = 0.883), as well as for the night between the 1^st^ and the 2^nd^ day of testing (Welch’s ANOVA: *F*_(4, 27.9)_ = 1.25, *p* = 0.312).

### Learning session

Participants exhibited a fast initial improvement on motor performance on the FTT during the learning session (see Fig. 2). The mixed-design ANOVA performed on the GPI revealed a main Block effect (*F*_(6.05, 344.61)_ = 15.40, *p* < 0.001, *η*^*2*^_p_ = 0.213), but no significant main Group effect (*F*_(4, 57)_ = 0.624, *p* = 0.647, *η*^*2*^_p_ = 0.042) or Block by Group interaction (*F*_(24.18, 344.61)_ = 1.50, *p* = 0.063, *η*^*2*^_p_ = 0.095). Hence, all Group conditions similarly improved throughout the learning session.

**Fig 2.**
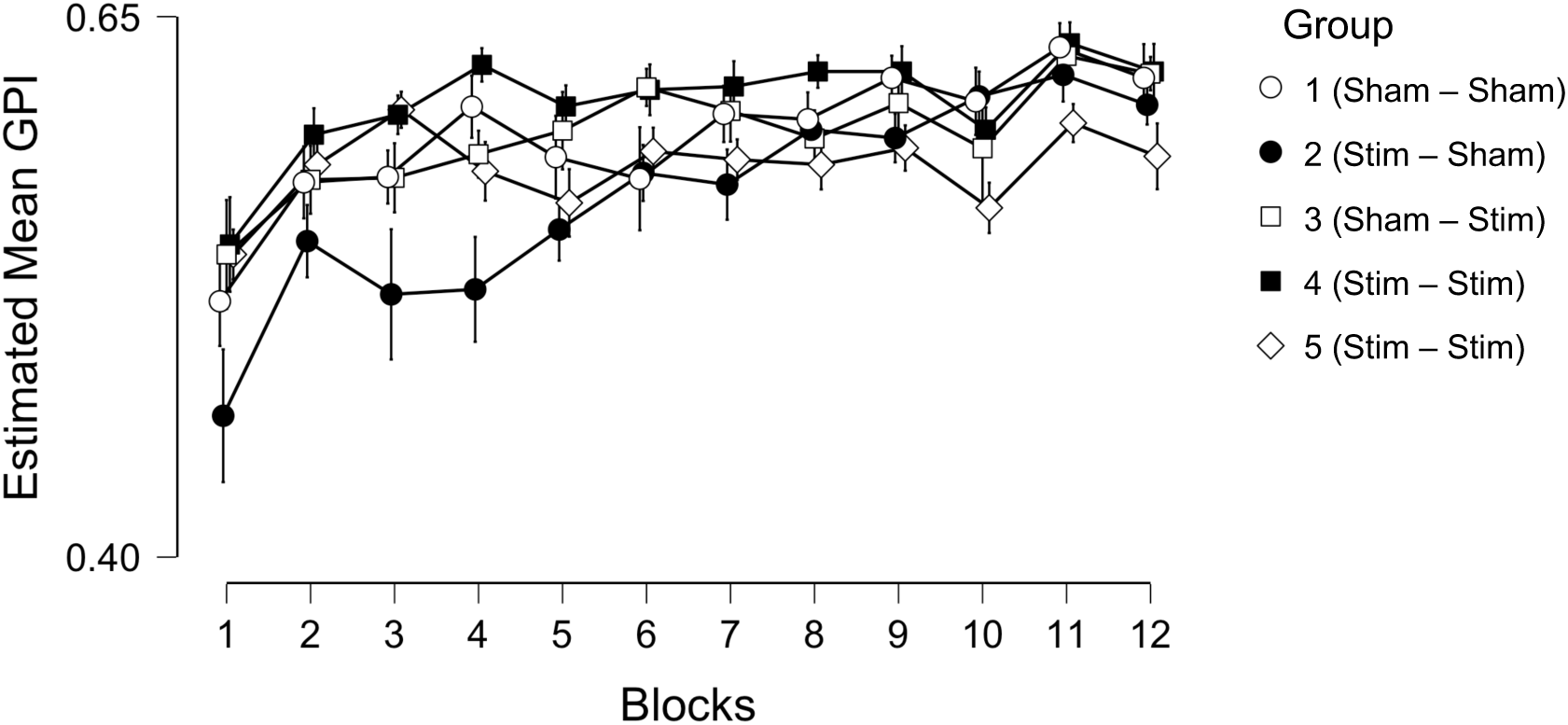
Evolution of global performance index (GPI) during the learning session (LS): Mean GPI and standard error over the 12 learning blocks for the 5 different Group conditions.

### Offline evolution of performance

Performance markedly increased from the end of the learning session (LS) to Test1, then stabilized between Test1 and Test2, to increase again at Test3 the next day, in a seemingly similar manner in the 5 Groups (Figure 3). Accordingly, the mixed-design ANOVA performed on GPI revealed a main Session effect (*F*_(2.49, 142.14)_ = 81.262, *p* < 0.001, *η*^*2*^_p_ = 0.588), but no significant main Group effect (*F*_(4, 57)_ = 0.282, *p* = 0.888, *η*^*2*^_p_ = 0.019) or Session by Group interaction (*F*_(9.98, 142.14)_ = 1.320, *p* = 0.225, *η*^*2*^_p_ = 0.085). Post-hoc pairwise comparisons decomposing the Session effect showed that all sessions significantly differed from each other (all *p*s < 0.001), but for Test1 vs. Test2 (*p* = 0.591).

**Fig 3.**
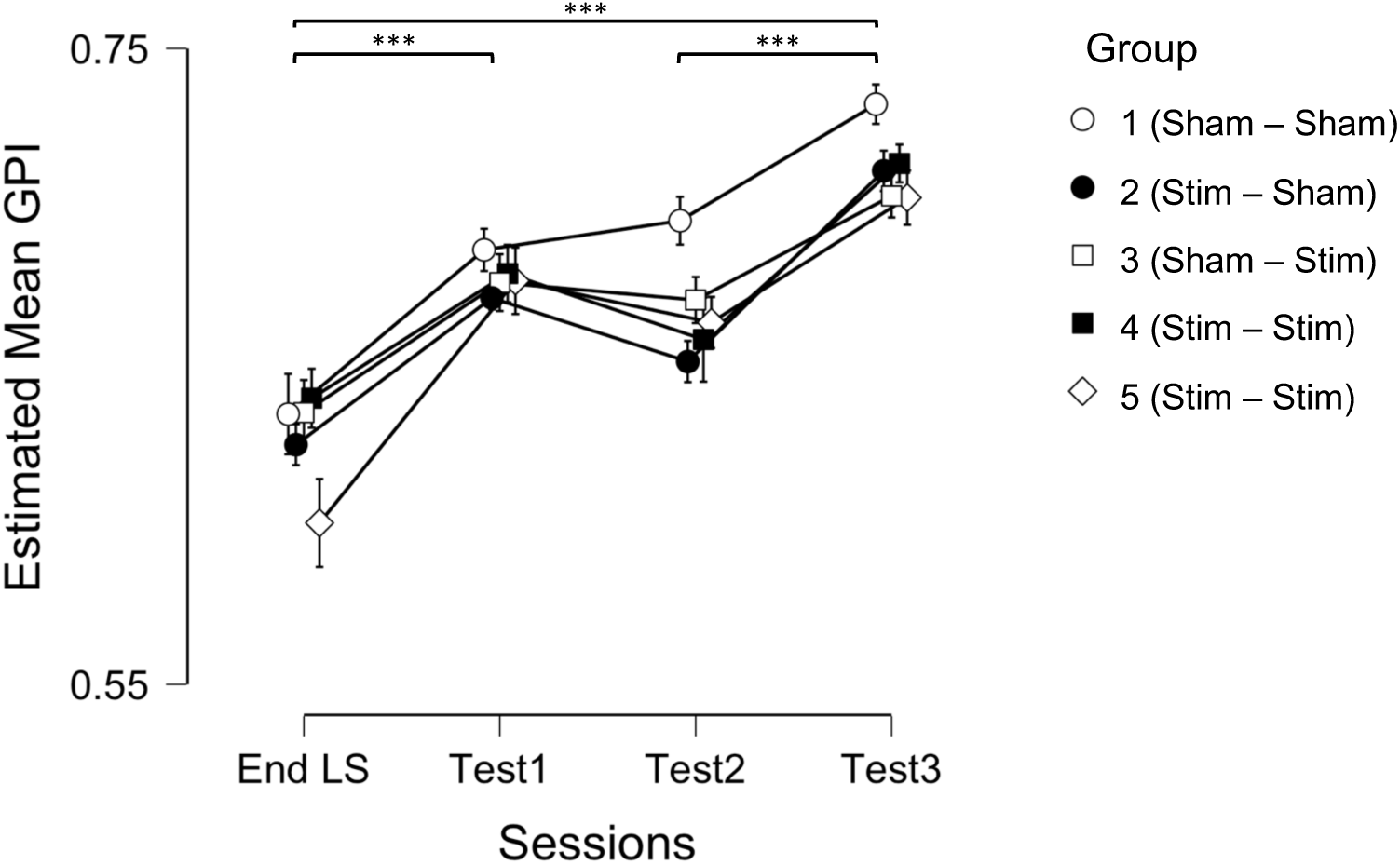
Global performance index (GPI) throughout sessions: Mean GPI and standard error across the end of learning (LS) and the 3 testing sessions held 25 min (Test 1), 4h (Test 2) and 24h (Test 3) post-learning for the 5 different Group conditions.

## Discussion

The present study aimed at investigating the effect of 20 Hz tACS on the evolution of motor performance within critical time windows for the expression of performance. All groups exhibited a similar evolution in performance over time, independently from the type of stimulation (20 Hz tACS or Sham tACS) received at the different time points.

In the present study, all conditions followed the motor learning framework described in prior reports [2], [4], with a significant increase in performance during the early boost period at 25 min, then a stabilization of performance gains within the silent period (4h) and a substantial improvement 24h later. However, none of our stimulation conditions significantly modulated memory consolidation performance, which is partially discrepant with previous reports. In particular, Krause et al. [18] evidenced a significant performance improvement after the application of 20 Hz tACS over M1 within the 10 minutes following practice a sequential motor task. Performance (i.e., speed) was measured at baseline, the end of acquisition, directly after stimulation and after a reacquisition session (identical to the initial training session) following the stimulation period. Their results highlighted enhanced speed immediately after the stimulation period when stimulating offline for 10 minutes in the 20 Hz tACS but not in the Sham tACS condition. Additionally, speed after reacquisition was similar in the 20 Hz and Sham tACS conditions, suggesting that 20 Hz tACS improved reaction times mostly in restricted training conditions. For the sake of comparison with Krause et al.’s [18] findings, we computed a similar analysis contrasting 20 Hz and Sham tACS conditions on speed values rather than the GPI. In line with Krause et al.’s results, our analysis showed that even if speed improved in both conditions, the amplitude of the improvement was higher (p < .05, see SI results) after 20 Hz than Sham tACS. However, this effect was not replicated when using the GPI parameter as performance measure (see SI results). This questions possible comparisons between studies using different parameters to reflect performance improvement and motor sequence learning. Indeed, speed and accuracy are partially subtended by different neural structures and seem to evolve relatively independently from each other [32], [33]. Furthermore, speed measures do not always adequality reflect subsequent cerebral plasticity [34] and, accordingly, the changes in memory. In this respect, combining speed and accuracy may provide a more reliable assessment of motor sequence learning.

This together with other methodological differences may explain some discrepancies between our and others’ reports. One important issue is the choice of the appropriate tACS montage. Several tACS studies used a cephalic montage with the active electrode located on M1 and the reference located above the contralateral orbital frontal area (Fo), whereas we used an extracephalic montage (M1 - contralateral deltoid; see SI Fig. 1). The extracephalic montage has the advantage of generating a negligible amount of phosphenes as compared to a M1-Fo montage [35], which makes it more easy to keep the participants blind about the type of stimulation they receive. This was confirmed by our participants who provided similar subjective reports about their sensations during the stimulation period, either actual or sham (see SI results). Furthermore, the extracephalic montage enables stimulating a narrower area by inducing larger vertical current densities in the somatosensory and primary motor cortex when located on M1 compared to cephalic montages (see SI Fig. 2). However, the horizontal current density is weaker but more homogeneous among the cortex. On the contrary, cephalic montages produce a larger current density on the cortical surface but more heterogeneously distributed [36]. Additionally, a contralateral Fo-M1 montage generates the strongest observable electric field in the frontal regions [32-34]. Likewise, for most individuals, a high electric field intensity is usually found in premotor areas, the fundus of the central sulcus, the postcentral gyrus, the postcentral sulcus [40], whereas high current flows are observed in the insula, cingulate cortex and the thalamus [38]. The incidental stimulation of these areas could play a role in the observed aftereffects. On the other hand, studies using extracephalic montages revealed discrepancies in the stimulation of deep brain structures. Some studies reported a lack of significant differences in the electric field among subcortical nuclei [39], while others revealed that the brainstem, cerebellum or basal ganglia might be stimulated by the large current spread [35]. Notably, whilst some areas are found to be consistently activated by stimulation among participants [35, 36], other regions seem to be differently affected due to anatomical variations in cortex shape [40], cerebrospinal fluid density [41] and skull thickness [42]. To sum up, cephalic montages essentially result in a heterogeneous stimulation of the cortex with a strong current density over frontal regions, whereas extracephalic montages tend to activate more specifically the target area with a weaker but homogeneous impact on the entire cortex, at the risk of deeper brain structures activation. There is also evidence that the magnitude of the elicited effect tends to decrease as the interelectrode distance increases [43], which is the case with an extracephalic montage. Thus, stimulating at a higher intensity might have been necessary to generate stronger aftereffects in our experiment.

While some electrical stimulation studies reported a high inter- and intra-individual reliability among sessions [44], others evidenced a large inter- and intraindividual variability [40-43]. Furthermore, in an experiment led with continuous theta-burst stimulation (cTBS) applied over M1 with TMS, only 50% of the participants responded to stimulation by showing decreased excitability [49]. To the best of our knowledge, this question has not been investigated yet using tACS. Nonetheless, we cannot rule out the possibility that inter-individual variability played a role in the apparent lack of effect of electrical stimulation in the present study. Finally, recent evidence suggests a crucial role of the stimulation phase in the elicited aftereffects. Previous studies showed no changes in motor evoked potentials (MEPs) when stimulating M1 with 20 Hz tACS without controlling for the phase [50], while increased MEPs are usually found at this frequency [12, 46, 47]. However, it has been shown that modulations of MEPs with β-tACS were phase-dependent in a way that only 20 Hz tACS at 90° succeeded to increase M1 excitability [53]. In our study, the stimulation was applied without the phase shift (0° phase) which may explain no substantive aftereffects observed.

To conclude, a large number of interconnected factors can influence the way tACS modulates brain activity. The modification of a single one (e.g. current intensity/density, electrodes placement, phase, stimulation duration, inter-individual and intra-individual differences, etc.) could impact the tACS outcomes. Furthermore, brain stimulation research is often associated with small to moderate effect sizes. The small sample sizes used in most studies could have led to an over- or underestimation of the impact of stimulation techniques [54]. At this stage, our data do not provide substantial support for a positive effect of the 20 Hz tACS on motor memory consolidation even when considering aggregated data contrasting 20 Hz tACS vs. Sham conditions (see SI). However, considering the increasing number of β-oscillations applications in rehabilitation therapy, further research remains needed to elucidate the capacity of 20 Hz tACS to modulate β-power and generate measurable aftereffects on motor function in humans.

## Acknowledgements

We are grateful to Tommaso H. Faret for his help in data acquisition.

## Author Contributions

LR and WS were the leading researchers for this study and contributed to all steps. PP contributed to the experimental design, data analysis and to the redaction of the article.

## Supplementary Information

### 1. Demographic information

**SI Table 1.**
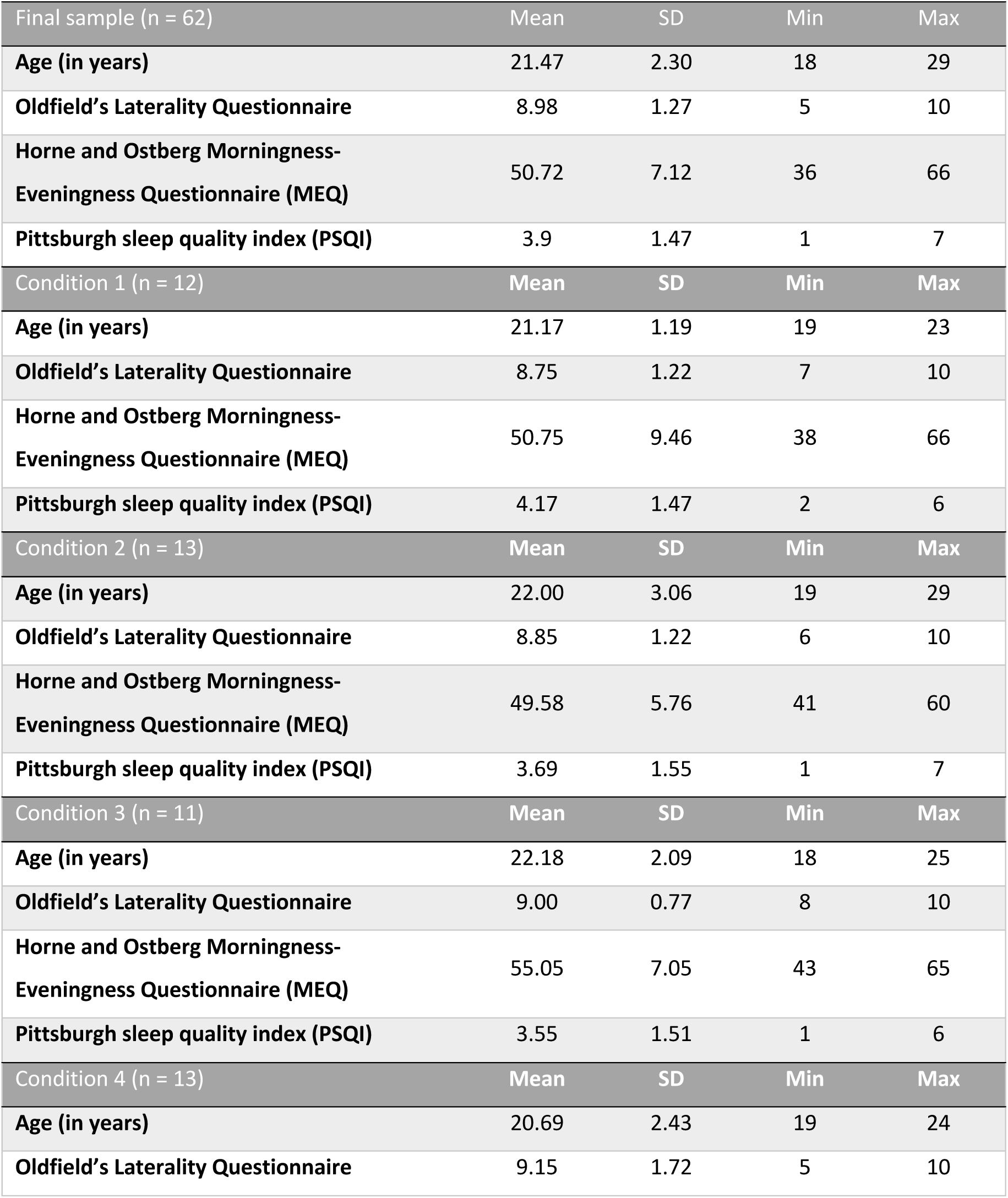
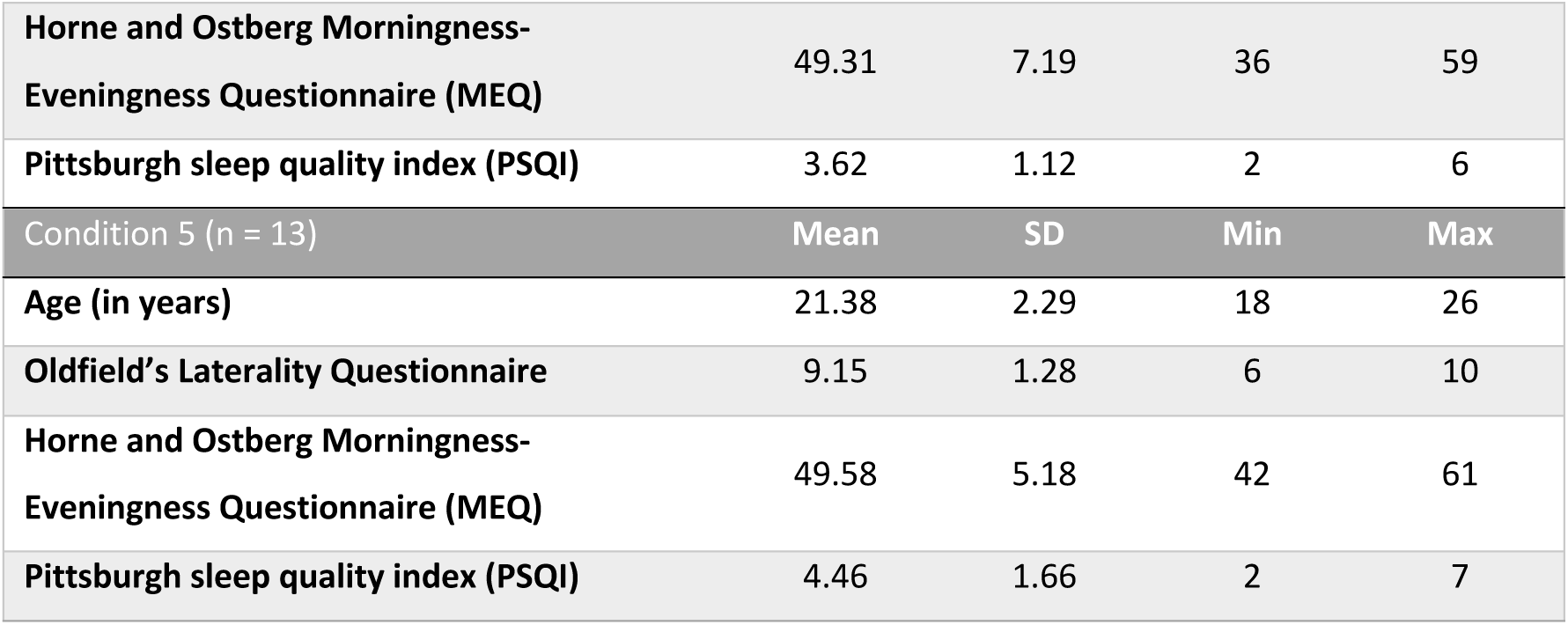
Table resuming mean (Mean), standard deviation (SD), minimum (Min) and maximum value (Max) for the total sample as well as for each of the 5 conditions (see main text) separately concerning age, laterality (Oldfield’s Laterality Questionnaire), circadian rhythm (Horne and Ostberg Morningness-Eveningness Questionnaire (MEQ)) and sleep quality (Pittsburgh sleep quality index (PSQI)).

| Final sample (n = 62) | Mean | SD | Min | Max |
| --- | --- | --- | --- | --- |
| Age (in years) | 21.47 | 2.30 | 18 | 29 |
| Oldfield's Laterality Questionnaire | 8.98 | 1.27 | 5 | 10 |
| Horne and Ostberg Morningness-Eveningness Questionnaire (MEQ) | 50.72 | 7.12 | 36 | 66 |
| Pittsburgh sleep quality index (PSQI) | 3.9 | 1.47 | 1 | 7 |
| Condition 1 (n = 12) | Mean | SD | Min | Max |
| Age (in years) | 21.17 | 1.19 | 19 | 23 |
| Oldfield's Laterality Questionnaire | 8.75 | 1.22 | 7 | 10 |
| Horne and Ostberg Morningness-Eveningness Questionnaire (MEQ) | 50.75 | 9.46 | 38 | 66 |
| Pittsburgh sleep quality index (PSQI) | 4.17 | 1.47 | 2 | 6 |
| Condition 2 (n = 13) | Mean | SD | Min | Max |
| Age (in years) | 22.00 | 3.06 | 19 | 29 |
| Oldfield's Laterality Questionnaire | 8.85 | 1.22 | 6 | 10 |
| Horne and Ostberg Morningness-Eveningness Questionnaire (MEQ) | 49.58 | 5.76 | 41 | 60 |
| Pittsburgh sleep quality index (PSQI) | 3.69 | 1.55 | 1 | 7 |
| Condition 3 (n = 11) | Mean | SD | Min | Max |
| Age (in years) | 22.18 | 2.09 | 18 | 25 |
| Oldfield's Laterality Questionnaire | 9.00 | 0.77 | 8 | 10 |
| Horne and Ostberg Morningness-Eveningness Questionnaire (MEQ) | 55.05 | 7.05 | 43 | 65 |
| Pittsburgh sleep quality index (PSQI) | 3.55 | 1.51 | 1 | 6 |
| Condition 4 (n = 13) | Mean | SD | Min | Max |
| Age (in years) | 20.69 | 2.43 | 19 | 24 |
| Oldfield's Laterality Questionnaire | 9.15 | 1.72 | 5 | 10 |
| Horne and Ostberg Morningness-Eveningness Questionnaire (MEQ) | 49.31 | 7.19 | 36 | 59 |
| Pittsburgh sleep quality index (PSQI) | 3.62 | 1.12 | 2 | 6 |
| Condition 5 (n = 13) | Mean | SD | Min | Max |
| Age (in years) | 21.38 | 2.29 | 18 | 26 |
| Oldfield's Laterality Questionnaire | 9.15 | 1.28 | 6 | 10 |
| Horne and Ostberg Morningness-Eveningness Questionnaire (MEQ) | 49.58 | 5.18 | 42 | 61 |
| Pittsburgh sleep quality index (PSQI) | 4.46 | 1.66 | 2 | 7 |

### 2. tACS parameters and subjective sensation reports

Cephalic vs extracephalic montages and related current density are illustrated SI Fig. 1 and SI Fig. 2, respectively.

For each stimulation period, impedance was kept below 10 kOhm. Mean impedance across stimulations was 6.75 ± 2.36 kOhm (min = 1.2 kOhm, max = 9.92 kOhm) and did not significantly differ between groups (Welch’s ANOVA: *F*_(4, 27.87)_ = 0.903, *p* = 0.476).

Mann-Whitney’s tests (Sham tACS vs. tACS Stimulation) separately performed on each type of sensation (auditory, visual, cutaneous or gustatory) reported by the subjects after the first and the second stimulation period did not disclose any significant difference (*p*s > 0.117). Hence, participants do not seem to have experienced different sensations regardless of the type of stimulation (Sham tACS vs. tACS Stimulation) received.

**SI Fig 1.**
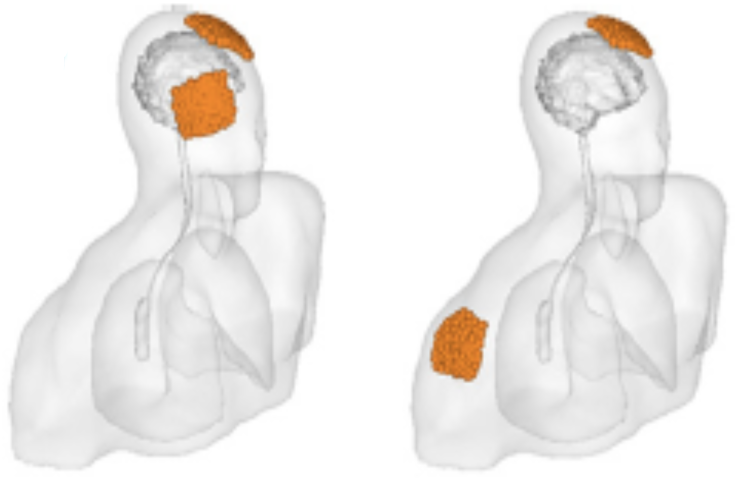
Cephalic montage (left) with active electrode on M1 and reference electrode placed on the contralateral supraorbital region (Fo), and extracephalic montage (right; our current experiment) with active electrode on M1 and reference electrode placed on the contralateral deltoid [1].

**SI Fig 2.**
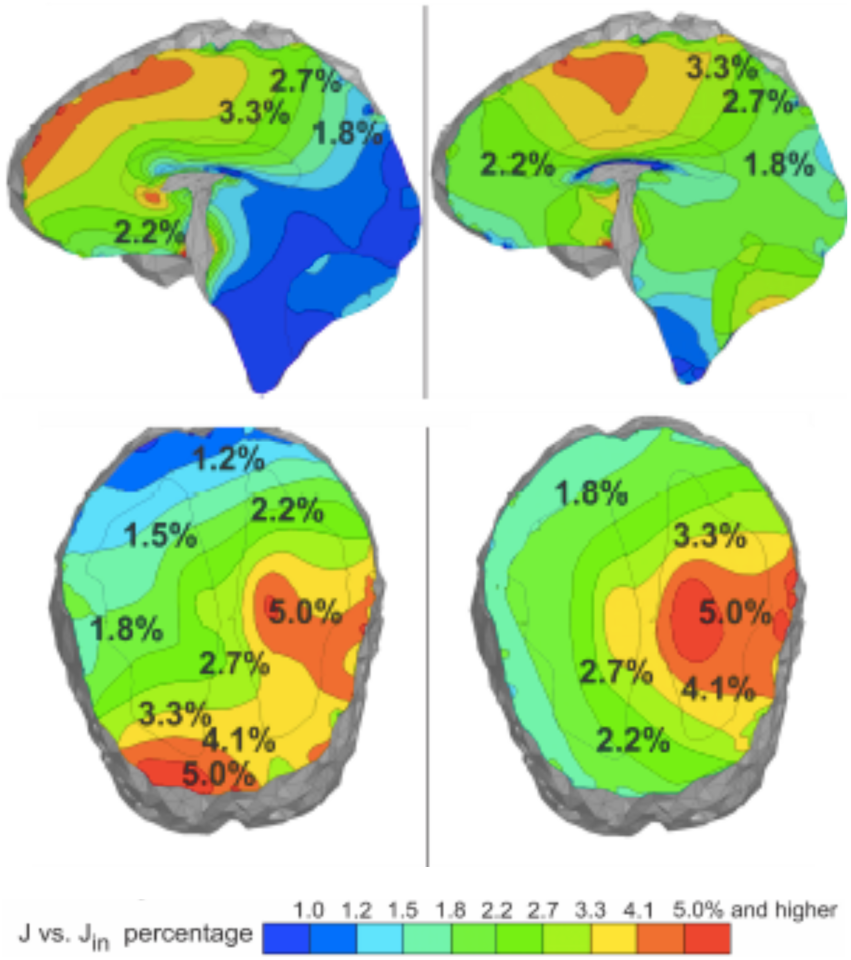
Computed total current density (J) normalized by the applied current density (J_in_) using a logarithmic scale for respectively the cephalic montage M1-contralateral Fo (left) and the extracephalic montage M1 – contralateral deltoid (right; our current experiment) [2].

### 3. Additional measures results (alertness, sleep and fatigue)

#### Alertness (PVT)

The mixed-design ANOVA conducted on the RRT [3] disclosed a main Session effect (*F*_(1.64, 93.49)_ = 8.433, *p* = 0.001). The Group (*p* = 0.646) and Group by Session interaction (*p =* 0.783) effects were non-significant, indicating that vigilance similarly evolved in the 5 Group conditions (see SI Figure 3). Post-hoc comparisons disclosed a significant difference between LS and Test 3 (*p* = 0.003), alertness being lower before the LS than before Test 3. A marginal difference was observed between LS and Test 2 (*p* = 0.053), but not between Test 2 and Test 3 (*p* = 0.123).

**SI Fig 3.**
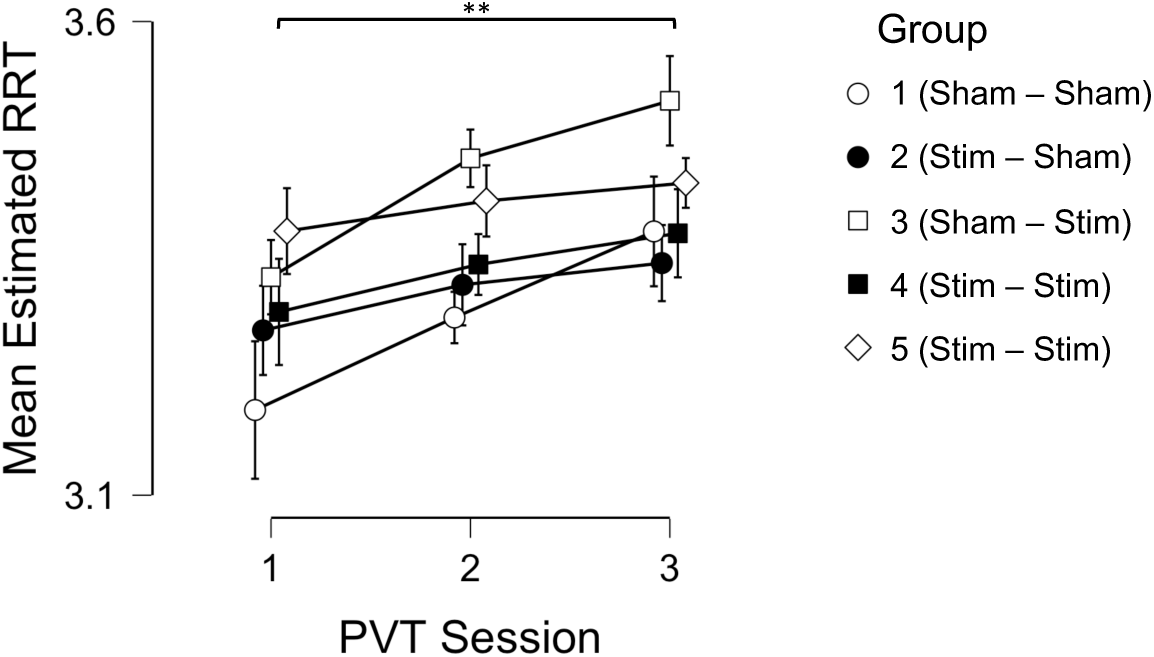
Alertness. Mean reciprocal reaction time (RRT) and standard error before learning (LS) and before testing sessions held 4h (Test2) and 24h (Test3) post-learning for the 5 Group conditions.

#### Sleepiness

The mixed-design ANOVA performed on Sleepiness with within-subject factors Time (Pre vs. Post), Session (LS, Test 1, Test 2, Test3) and between-subject factor Group (1-5) revealed main effects of Time (*F*_(1, 57)_ = 26.954, *p* < 0.001) and Session (*F*_(2.60, 148.39)_ = 4.861, *p* = 0.005). The Group (*p* = 0.904), Time by Session (*p* = 0.946), Time by Group (*p* = 0.051), Session by Group (*p* = 0.601) and Time by Session by Group interaction (*p =* 0.767) effects were non-significant, suggesting that sleepiness similarly evolved in the 5 Group conditions (see SI Figure 4). Post-hoc comparisons disclosed significant Pre vs. Post difference, sleepiness being higher after the task as compared as before. Post-hoc comparisons on the Session effect disclosed a higher sleepiness at Test 3 than LS (*p* = 0.003). No other significant difference was found across the different sessions (all *p*s > 0.067).

**SI Fig 4.**
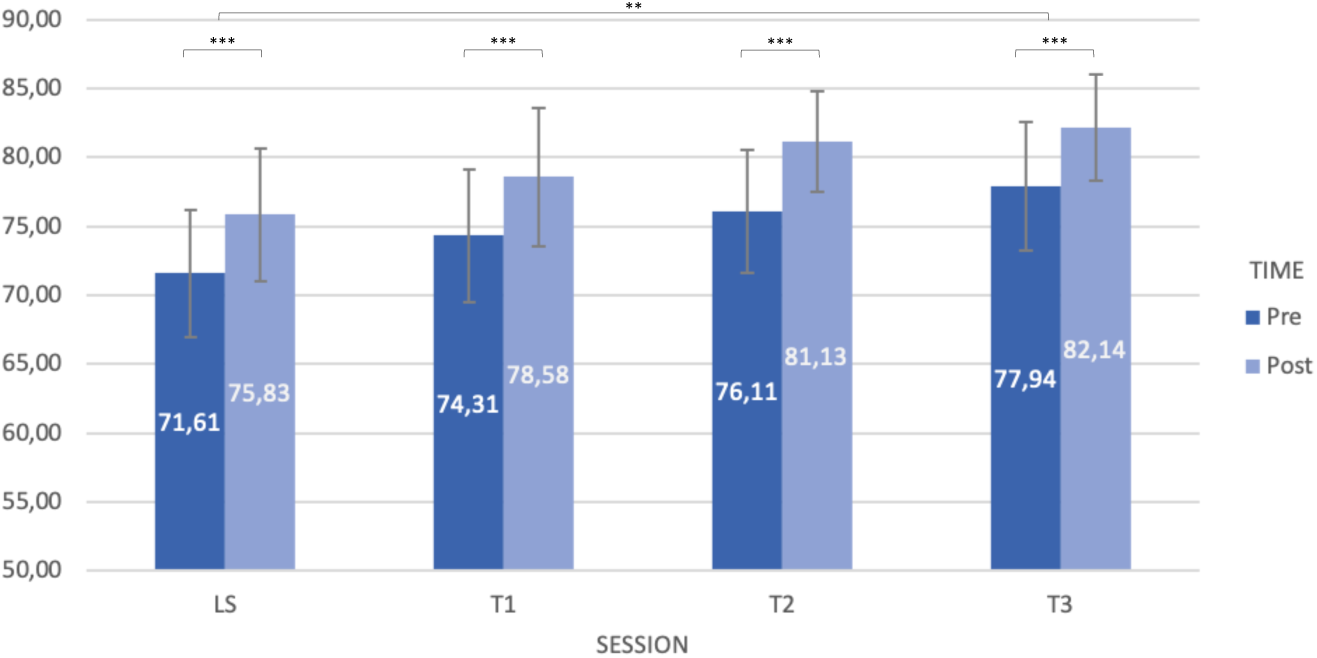
Sleepiness. Mean sleepiness % and standard error before and after (Pre vs. Post) the learning session (LS) and the 3 testing sessions held 25 min (Test1), 4h (Test2) and 24h (Test3) post-learning. The higher the score, the higher the subjective feeling of sleepiness.

#### Fatigue

The mixed-design ANOVA conducted on Fatigue with within-subject factors Time (Pre vs. Post) and Session (LS, Test 1, Test 2, Test3) and between-subjects Group factor (1-5) revealed a main effect of Time (*F*_(1, 57)_ = 1.767, *p* = 0.001) and a Time by Session interaction (*F*_(2.64, 150.63)_ = 4.725, *p* = 0.005). Main effect of Session (*p* = 0.092), Group (*p* = 0.629), as well as Time by Group (*p* = 0.608), Session by Group (*p* = 0.926) and Time by Session by Group interaction effects (*p =* 0.645) were non-significant. Post-hoc comparisons disclosed significant difference between Pre and Post (*p* < 0.001) ratings (see SI Figure 5), with a higher fatigue level after the task than before, but for LS, which explained the Time by Session interaction effect.

**SI Fig 5.**
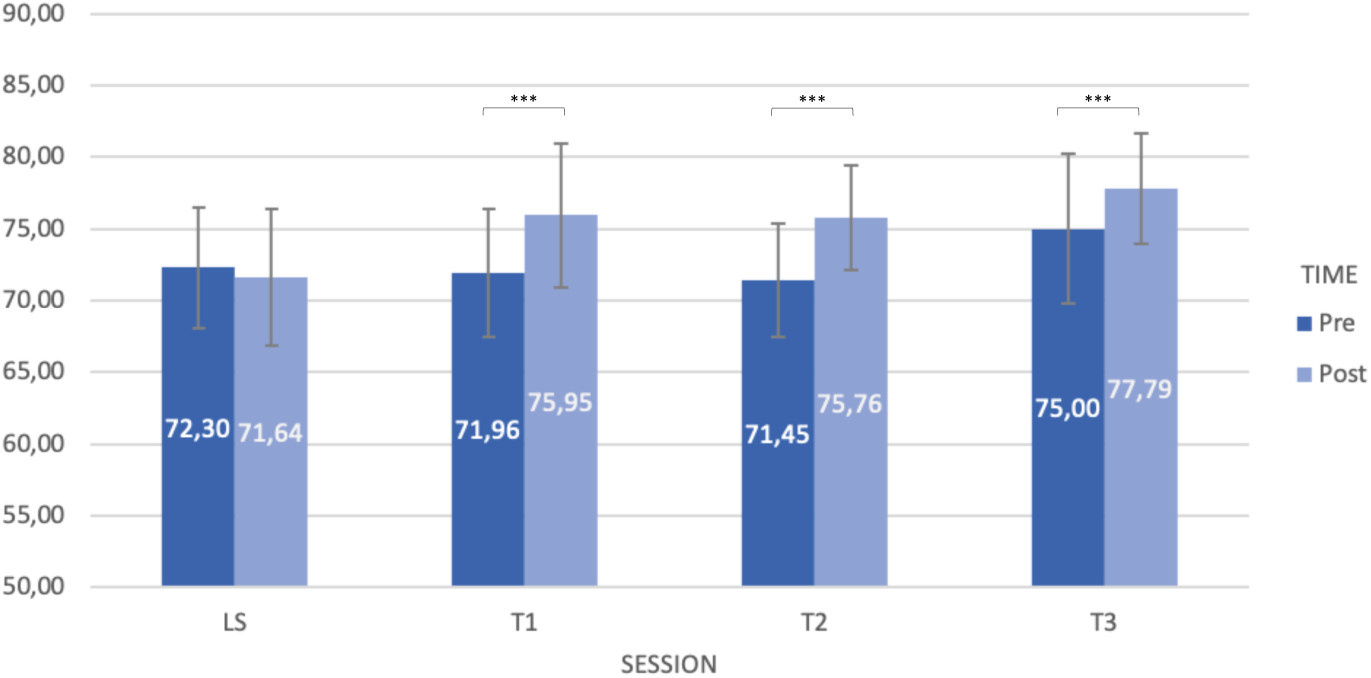
Fatigue. Mean fatigue % and standard error before and after (Pre vs. Post) the learning session (LS) and the 3 testing sessions held 25 min (Test1), 4h (Test2) and 24h (Test3) post-learning. The higher the score, the higher the subjective feeling of fatigue.

### 4. Additional FTT analyses

#### Offline evolution of performance (speed parameter)

For the purpose of comparisons with prior reports, we computed the offline evolution of performance using the speed parameter only (i.e., mean RT per block for correct responses) averaged over the two blocks of each test (Tests 1-3) as well as the last two blocks of the learning session (LS; see SI Figure 6). The mixed-design ANOVA on mean RTs evidenced a main Session effect (*F*_(1.98, 113.08)_ = 90.96, *p* < 0.001, *η*^*2*^_p_ = 0.615). The Group (*F*_(4, 57)_ = 0.726, *p* = 0.578, *η*^*2*^_p_ = 0.048) and Group by Session interaction (*F*_(7.94, 113.08)_ = 1.72, *p* = 0.102, *η*^*2*^_p_ = 0.010) effects were non-significant. Post-hoc pairwise comparisons decomposing the Session effect showed that reaction time decreased over sessions (all *p*s < 0.001) but for Test1 vs. Test2 (*p* = 0.518), replicating the results obtained using the GPI as a measure of performance (see main text results).

**SI Fig 6.**
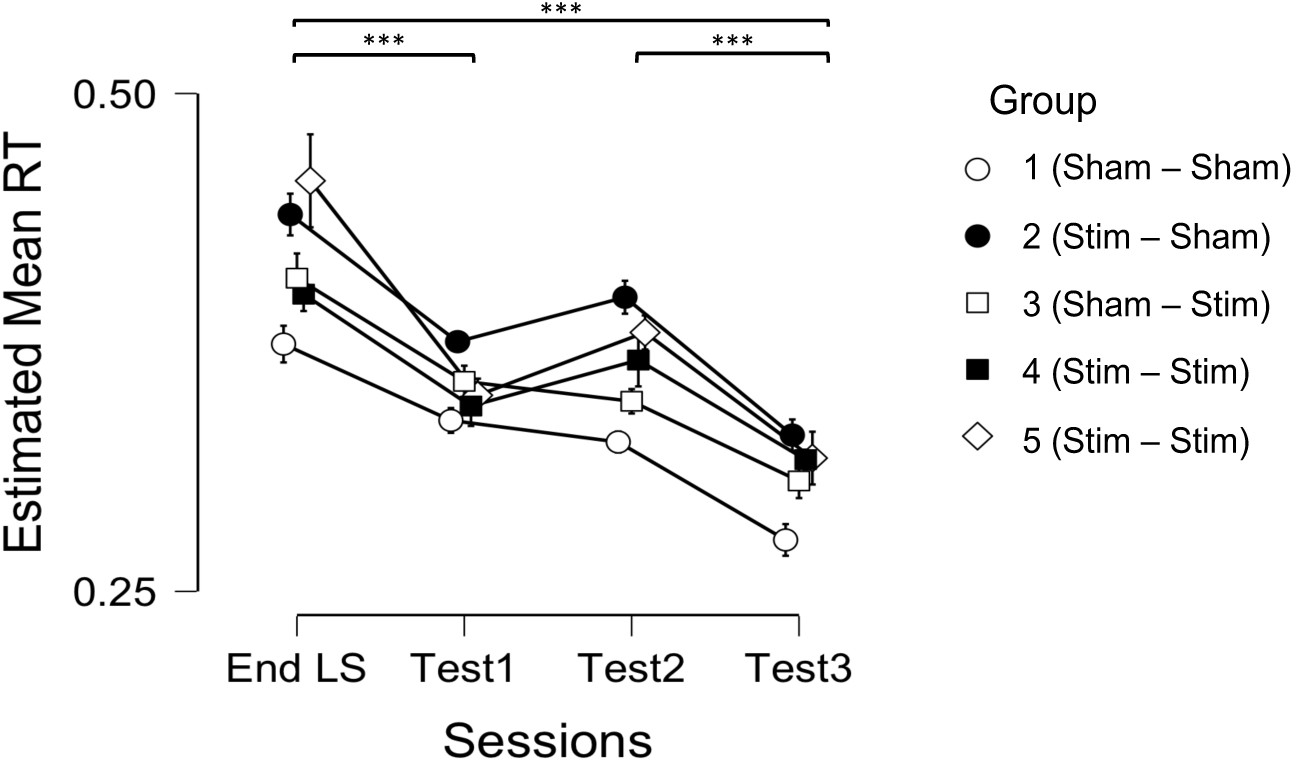
Offline speed evolution. Mean reaction time (RT) and standard error at the end of the learning session (End LS) and the 3 testing sessions held 25 min (Test1), 4h (Test2) and 24h (Test3) post-learning for the 5 different Group conditions.

#### Short-term effects of 20 Hz tACS vs Sham on performance

For the sake of comparison with Krause et al. (2016) [4] results, we computed an analysis contrasting 20 Hz vs. Sham tACS conditions on speed (mean RTs; see SI Figure 7). A mixed-design ANOVA investigated the evolution of speed before vs. after the stimulation period depending on the type of stimulation received (Sham vs. effective tACS). Thirty-nine participants were stimulated with 20 HZ tACS for 10minn whereas 23 received Sham tACS. The ANOVA revealed a main Session effect (*F*_(1, 60)_ = 60.75, *p* < 0.001, *η*^*2*^_p_ = 0.503) with a significant reaction time improvement between the end of the learning session (LS) and Test 1 (held 25min later), and a Stimulation by Session interaction effect (*F*_(1, 60)_ = 4.04, *p* = 0.049, *η*^*2*^_p_ = 0.063). The main Stimulation effect (*F*_(1, 60)_ = 1.02, *p* = 0.317, *η*^*2*^_p_ = 0.017) was non-significant. Post-hoc independent t-tests decomposing the Stimulation by Session effect did not evidence differences between the 2 stimulation conditions (Sham vs. effective tACS) at LS (t_(60)_ = -1.374; *p* = 0.174) or at Test 1 (t_(60)_ = -0.452; *p* = 0.653). T-test for dependent samples evidenced a significant reaction time improvement both in the Sham (t_(22)_ = 4.89; *p* < 0.001) and effective tACS (t_(38)_ = 7.18; *p* < 0.001) conditions.

**SI Fig 7.**
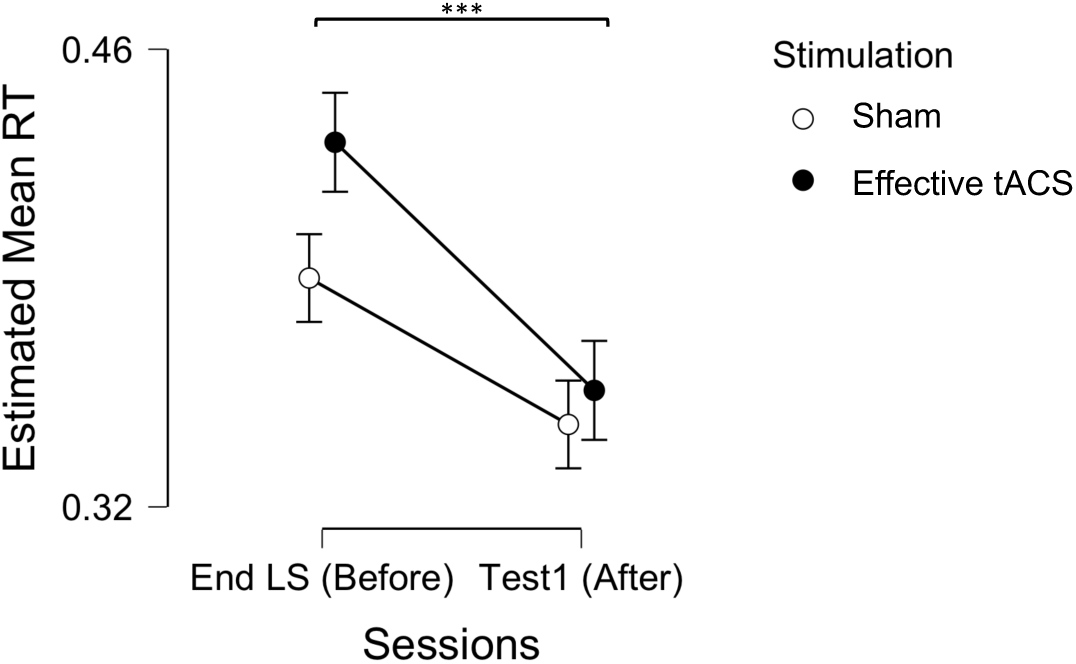
Evolution of speed (RTs) after Sham vs. effective tACS. Mean reaction time (RT) and standard error at the end of the learning session (End LS) and the testing session held after 25 min (Test1).

To explore further the source of the Stimulation by Session interaction effect, we computed the delta (difference) between RTs at LS and Test 1 in the tACS and Sham conditions. A t-test for independent samples revealed that performance improvement was more pronounced in the tACS than the Sham condition (t_(60)_ = -2.01; *p* = 0.049). Taken together, these results suggest that our interaction effect was driven by a stronger speed improvement in the effective tACS (mean difference = 0.0759) than Sham (mean difference = 0.0448) condition.

Finally and for the sake of completeness in our comparisons, we computed a similar mixed design ANOVA on the GPI (see SI Figure 8). This analysis revealed a main Session effect (F_(1, 60)_ = 56.849, *p* < 0.001, η^2^_p_ = 0.487) with a significant GPI increase between the end of LS and Test 1 (held 25min later). However, the main Stimulation (F_(1, 60)_ = 0.300, *p* = 0.586, η^2^_p_ = 0.005) and the Stimulation by Session interaction (F_(1, 60)_ = 0.342, *p* = 0.561, η^2^_p_ = 0.006) effects were non-significant.

**SI Fig 8.**
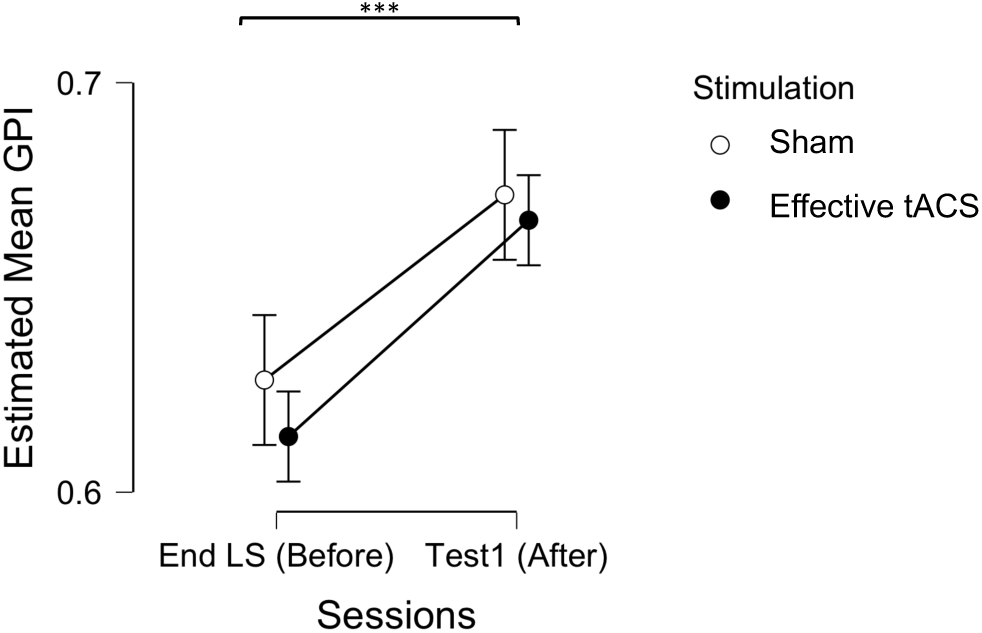
Evolution of GPI after Sham vs. effective tACS. Mean reaction time (RT) and standard error at the end of the learning session (End LS) and the testing session held after 25 min (Test1).

## References

[1] Karni A, Meyer G, Rey-Hipolito C, Jezzard P, Adams M, Turner R, Ungerleider L. The acquisition of skilled motor performance: Fast and slow experience-driven changes in primary motor cortex. Proc Natl Acad Sci 1998;95(3):861–868. https://doi.org/10.1073/pnas.95.3.861.

[2] Hotermans C, Peigneux P, De Noordhout A. M, Moonen G, Maquet P. Early boost and slow consolidation in motor skill learning. Learn Mem 2006;13:580–583. https://doi.org/10.1101/lm.239406.

[3] Hotermans C, Peigneux P, De Noordhout AM, Moonen G, Maquet P. Repetitive transcranial magnetic stimulation over the primary motor cortex disrupts early boost but not delayed gains in performance in motor sequence learning. Eur J Neurosci 2008;28(6):1216–1221. https://doi.org/10.1111/j.1460-9568.2008.06421.x.

[4] Albouy G, Ruby P, Phillips C, Luxen A, Peigneux P, Maquet P. Implicit oculomotor sequence learning in humans: Time course of offline processing. Brain Res 2006;1090(1):163–71. https://doi.org/10.1016/j.brainres.2006.03.076.

[5] Schmitz R, Schabus M, Perrin F, Luxen A, Maquet P, Peigneux P. Recurrent boosting effects of short inactivity delays on performance: An ERPs study. BMC Res Notes 2009;2(1):170. https://doi.org/10.1186/1756-0500-2-170.

[6] Borragán G, Urbain C, Schmitz R, Mary A, Peigneux P. Sleep and memory consolidation: Motor performance and proactive interference effects in sequence learning. Brain Cogn 2015;95:54–61. https://doi.org/10.1016/j.bandc.2015.01.011.

[7] Halsband U, Lange RK. Motor learning in man: A review of functional and clinical studies. J Physiol Paris 2006;99(4-6):414–24. https://doi.org/10.1016/j.jphysparis.2006.03.007.

[8] Sami S, Robertson EM, Miall RC. The time course of task-specific memory consolidation effects in resting state networks. J Neurosci 2014;34(11):3982–3992. https://doi.org/10.1523/JNEUROSCI.4341-13.2014.

[9] Pollok B, Latz D, Krause V, Butz M, Schnitzler A. Changes of motor-cortical oscillations associated with motor learning. Neuroscience 2014;275(5):47–53. https://doi.org/10.1016/j.neuroscience.2014.06.008.

[10] Thut G, Miniussi C, Gross J, Miniussi C, Gross J. The functional importance of rhythmic activity in the brain. Curr Biol 2012;22(16):R658–R663. https://doi.org/10.1016/j.cub.2012.06.061.

[11] Helfrich RF, Schneider TR, Rach S, Trautmann-Lengsfeld SA, Engel AK, Herrmann CS. Entrainment of brain oscillations by transcranial alternating current stimulation. Curr Biol 2014;24(3):333–339. https://doi.org/10.1016/j.cub.2013.12.041.

[12] Veniero D, Vossen A, Gross J, Thut G, Quentin R. Lasting EEG / MEG aftereffects of rhythmic transcranial brain stimulation: level of control over oscillatory network activity. Front Cell Neurosci 2015;9:477. https://doi.org/10.3389/fncel.2015.00477.

[13] Kasten FH, Dowsett J, Herrmann CS. Sustained aftereffect of α - tACS lasts up to 70 min after stimulation. Front Hum Neurosci 2016;10:245. https://doi.org/10.3389/fnhum.2016.00245.

[14] Wischnewski M, Engelhardt M, Salehinejad MA, Kuo M, Nitsche MA, Schutter DJLG. NMDA receptor-mediated motor cortex plasticity after 20 Hz transcranial alternating current stimulation. Cereb Cortex 2019;29(7):2924–2931. https://doi.org/10.1093/cercor/bhy160.

[15] Feurra M, Bianco G, Santarnecchi G, Testa M, Rossi A, Rossi S. Frequency-dependent tuning of the human motor system induced by transcranial oscillatory potentials. J Neurosci 2011;31(34):12165–12170. https://doi.org/10.1523/JNEUROSCI.0978-11.2011.

[16] Weinrich CA, Brittain JS, Nowak M, Salimi-Khorshidi R, Brown P, Stagg CJ. Modulation of long-range connectivity patterns via frequency-specific stimulation of human cortex. Curr Biol 2017;(19):3061-3068.e3. https://doi.org/10.1016/j.cub.2017.08.075.

[17] Pollok B, Boysen A, Krause V. The effect of transcranial alternating current stimulation (tACS) at alpha and beta frequency on motor learning. Behav Brain Res 2015; 293:234–240. https://doi.org/10.1016/j.bbr.2015.07.049.

[18] Krause V, Meier V, Dinkelbach L, Pollok B. Beta band transcranial alternating (tACS) and direct current stimulation (tDCS) applied after initial learning facilitate retrieval of a motor sequence. Front Behav Neurosci 2016;10:1–10. https://doi.org/10.3389/fnbeh.2016.00004.

[19] Rumpf J, Barbu A, Fricke C, Wegscheider M, Classen C. Posttraining alpha transcranial alternating current stimulation impairs motor consolidation in elderly people. Neural Plast 2019;2019:ID2689790. https://doi.org/10.1155/2019/2689790.

[20] Horne JA, Ostberg O. A self-assessment questionnaire to determine morningness eveningness in human circadian rhythms. Int J Chronobiol 1976;4(2):97-110. PMID: 1027738.

[21] Buysse DJ, Reynolds CF, Monk TH, Berman SR, Kupfer DJ. The Pittsburgh sleep quality index: a new instrument psychiatric practice and research. Psychiatry Res 1989;28(2):193–213. https://doi.org/10.1016/0165-1781(89)90047-4.

[22] Genzel L, Kiefer T, Renner L, Wehrle R, Kluge M, Grözinger M, Steiger A, Dresler M. Sex and modulatory menstrual cycle effects on sleep related memory consolidation. Psychoneuroendocrinology 2012;37(7):987–998. https://doi.org/10.1016/j.psyneuen.2011.11.006.

[23] Renner L, Kiefer T, &, Genzel L, Dresler M, Pawlowski M, Steiger A. Menstrual cycle effects on sleep-dependent memory consolidation. Exp Clin Endocrinol Diabetes 2010;118–P17. DOI: 10.1055/s-0030-1267019.

[24] Klem GH, Lüders HO, Jasper HH, Elger C. The ten-twenty electrode system of the International Federation. The International Federation of Clinical Neurophysiology. Electroencephalography and Clinical neurophysiology. Supplement. 1999;52:3-6. PMID: 10590970.

[25] Rich, TL, Menk, JS, Rudser KD, Chen M, Meekins GD, Peña E, Feyma T, Bawroski K, Bush C, Gillick BT. Determining electrode placement for transcranial direct current stimulation: A comparison of EEG-versus TMS-guided methods. Clin EEG Neurosci 2017;48(6):367–375. https://doi.org/10.1177/1550059417709177.

[26] Ellis BW, Johns MW, Lancaster R, Raptopoulos P, Angelopoulos N, Priest NG. The St. Mary’s hospital sleep questionnaire: a study of reliability. Sleep 1981;4(1):93–97. https://doi.org/10.1093/sleep/4.1.93.

[27] Dinges DF, Powell JW. Microcomputer analyses of performance on a sustained operations. Behav Res Methods 1985;17(6):652–655. https://doi.org/10.3758/BF03200977.

[28] Basner M, Mcguire S, Goel N, Rao H, Dinges DF. A new likelihood ratio metric for the psychomotor vigilance test and its sensitivity to sleep loss. J Sleep Res 2015;24(6):702–713. https://doi.org/10.1111/jsr.12322.

[29] Lee KA, Hicks G, Nino-Murcia G. Validity and reliability of a scale to assess fatigue. Psychiatry Res 1991;36(3):291–298. https://doi.org/10.1016/0165-1781(91)90027-M.

[30] Laventure S, Fogel S, Lungu O, Albouy G, Sévigny-Dupont P, Vien C, Sayour C, Carrier J, Benali H, Doyon J. NREM2 and sleep spindles are instrumental to the consolidation of motor sequence memories. PLoS Biol 2016;31;14(3):e1002429. https://doi.org/10.1371/journal.pbio.1002429.

[31] Delacre M, Leys C, Mora YL, Lakens D. Taking parametric assumptions seriously: arguments for the use of Welch’s F-test instead of the Classical F-test in One-Way ANOVA. Int Rev Soc Psychol 2019;32(1):1–12. http://doi.org/10.5334/irsp.198.

[32] Adi-japha E, Fox O, Karni A. Research in developmental disabilities atypical acquisition and atypical expression of memory consolidation gains in a motor skill in young female adults with ADHD. Res Dev Disabil 2011;32(3):1011–1020. https://doi.org/10.1016/j.ridd.2011.01.048.

[33] Reitz S, Heisterübera M, Karni A, Gal C, Doyon J, King BR, Classen J, Rump J, Buccino G, Klann J, Binkofski F. EP 121. Motor sequence learning in patients with limb apraxia the effects of long-term training. Clin Neurophysiol 2016;127(9):e291–e292. https://doi.org/10.1016/j.clinph.2016.05.163.

[34] Doyon J, Orban P, Barakat M, Debas K, Lungu O, Albouy G, Fogel S, Proulx S, Laventure S, Deslauriers J, Duchesne C, Carrier J, Benali H. Plasticité fonctionnelle du cerveau et apprentissage moteur. Med Sci 2011;27(4):413–420. https://doi.org/10.1051/medsci/2011274018.

[35] Mehta AR, Pogosyan A, Brown P, Brittain JS. Montage matters: the influence of transcranial alternating current stimulation on human physiological tremor. Brain Stimul 2015;8(2):260–8. https://doi.org/10.1016/j.brs.2014.11.003.

[36] Noetscher GM, Member S, Yanamadala J, Makarov SN, Pascual-Leone A. Comparison of cephalic and extracephalic montages for transcranial direct current stimulation a numerical study. IEEE Trans Biomed Engy 2014;61(9):2488–98. DOI: 10.1109/TBME.2014.2322774.

[37] Datta A, Bansal V, Diaz J, Patel J, Reato D, Bikson M. Gyri-precise head model of transcranial direct current stimulation: Improved spatial focality using a ring electrode versus conventional rectangular pad. Brain Stimul 2009;2(4):201-207.e1. https://doi.org/10.1016/j.brs.2009.03.005.

[38] DaSilva AF, Truong DQ, DosSantos MF, Toback RL, Datta A, Bikson M. State-of-art neuroanatomical target analysis of high-definition and conventional tDCS montages used for migraine and pain control. Front Neuroanat 2015;9:1–12. https://doi.org/10.3389/fnana.2015.00089.

[39] Im CH, Park JH, Shim M, Chang WH, Kim YH. Evaluation of local electric fields generated by transcranial direct current stimulation with an extracephalic reference electrode based on realistic 3D body modelling. Phys Med Biol 2012;57(8):2137–2150. https://doi.org/10.1088/0031-9155/57/8/2137.

[40] Laakso I, Tanaka S, Mikkonen M, Koyama S, Sadato N, Hirata A. Electric fields of motor and frontal tDCS in a standard brain space: A computer simulation study. Neuroimage 2016;137:140–151. https://doi.org/10.1016/j.neuroimage.2016.05.032.

[41] Laakso I, Tanaka S, Koyama S, De Santis V, Hirata A. Inter-subject variability in electric fields of motor cortical tDCS. Brain Stimul 2015;8(5):906–913. https://doi.org/10.1016/j.brs.2015.05.002.

[42] Opitz A, Paulus W, Will S, Antunes A, Thielscher A. Determinants of the electric field during transcranial direct current stimulation. Neuroimage 2015;109:140–150. https://doi.org/10.1016/j.neuroimage.2015.01.033.

[43] Moliadze V, Antal A, Paulus W. Boosting brain excitability by transcranial high frequency stimulation in the ripple range. J Physiol 2010;24:4891–4904. https://doi.org/10.1113/jphysiol.2010.196998.

[44] Ammann C, Lindquist MA, Celnik PA. Response variability of different anodal transcranial direct current stimulation intensities across multiple sessions. Brain Stimul 2017;10(4):757–763. https://doi.org/10.1016/j.brs.2017.04.003.

[45] Chew T, Ho K, Loo CK. Inter- and Intra-individual variability in response to transcranial direct current stimulation (tDCS) at varying current intensities. Brain Stimul 2015;8(6):1130–1137. https://doi.org/10.1016/j.brs.2015.07.031.

[46] Dyke K, Kim S, Jackson GM, Jackson SR. Intra-subject consistency and reliability of response following 2 mA transcranial direct current stimulation. Brain Stimul 2016;9(6):819–825. https://doi.org/10.1016/j.brs.2016.06.052.

[47] López-Alonso V, Fernández-del-Olmo M, Costantini A, Gonzalez-Henriquez JJ, Cheeran B. Intra-individual variability in the response to anodal transcranial direct current stimulation. Clin Neurophysiol 2015;126:2342–2347. https://doi.org/10.1016/j.clinph.2015.03.022.

[48] Wiethoff S, Hamada M, Rothwell JC. Brain stimulation variability in response to transcranial direct current stimulation of the motor cortex. Brain Stimul 2014;7(3):468–475. https://doi.org/10.1016/j.brs.2014.02.003.

[49] Mcallister CJ, Ro KC, Stanford IM, Woodhall GL, Furlong PL, Hall SD. Oscillatory beta activity mediates neuroplastic effects of motor cortex stimulation in humans. J NEUROSCI 2013;33(18):7919–7927. https://doi.org/10.1523/JNEUROSCI.5624-12.2013.

[50] Guerra A, Pogosyan A, Nowak M, Tan H, Ferreri F, Di Lazzaro V, Brown P. Phase dependency of the human primary motor cortex and cholinergic inhibition cancelation during beta tACS. Cereb Cortex 2016;26(10):3977–90. https://doi.org/10.1093/cercor/bhw245.

[51] Cancelli A, Cottone C, Zito G, Di Giorgio M, Pasqualetti P, Tecchio F. Cortical inhibition and excitation by bilateral transcranial alternating current stimulation. Restor Neurol Neurosci 2015;33(2):105–114. https://doi.org/10.1016/j.clinph.2015.11.328.

[52] Feurra M, Blagovechtchenski E, Nikulin VV, Nazarova M, Lebedeva A, Pozdeeva D, Yurevich M, Rossi S. State-dependent effects of transcranial oscillatory currents on the motor system during action observation. Sci Rep 2019;9:12858. https://doi.org/10.1038/s41598-019-49166-1.

[53] Nakazono H, Ogata K, Kuroda T, Tobimatsu S. Phase and frequency-dependent effects of transcranial alternating current stimulation on motor cortical excitability. PLoS One 2016;11(9):e0162521. https://doi.org/10.1371/journal.pone.0162521.

[54] Veniero D, Benwell CYS, Ahrens MM, Thut G. Inconsistent effects of parietal α-tACS on pseudoneglect across two experiments: A failed internal replication. Front Psychol 2017;8:952. https://doi.org/10.3389/fpsyg.2017.00952.

## SI References

[1] Im CH, Park JH, Shim M, Chang WH, Kim YH. Evaluation of local electric fields generated by transcranial direct current stimulation with an extracephalic reference electrode based on realistic 3D body modelling. Phys Med Biol 2012;57(8):2137–2150. https://doi.org/10.1088/0031-9155/57/8/2137.

[2] Noetscher GM, Member S, Yanamadala J, Makarov SN, Pascual-Leone A. Comparison of cephalic and extracephalic montages for transcranial direct current stimulation a numerical study. IEEE Trans Biomed Engy 2014;61(9):2488–98. DOI: 10.1109/TBME.2014.2322774.

[3] Basner M, Mcguire S, Goel N, Rao H, Dinges DF. A new likelihood ratio metric for the psychomotor vigilance test and its sensitivity to sleep loss. J SLEEP RES 2015;24(6):702–713. https://doi.org/10.1111/jsr.12322.

[4] Krause V, Meier V, Dinkelbach L, Pollok B. Beta band transcranial alternating (tACS) and direct current stimulation (tDCS) applied after initial learning facilitate retrieval of a motor sequence. Front Behav Neurosci 2016;10:1–10. https://doi.org/10.3389/fnbeh.2016.00004.

